# Human Cytokine and Coronavirus Nucleocapsid Protein Interactivity Using Large-Scale Virtual Screens

**DOI:** 10.1101/2023.11.28.569056

**Authors:** Phillip J. Tomezsko, Colby T. Ford, Avery E. Meyer, Adam M. Michaleas, Rafael Jaimes

## Abstract

Understanding the interactions between SARS-CoV-2 and the human immune system is paramount to the characterization of novel variants as the virus co-evolves with the human host. In this study, we employed state-of-the-art molecular docking tools to conduct large-scale virtual screens, predicting the binding affinities between 64 human cytokines against 17 nucleocapsid proteins from six betacoronaviruses. Our comprehensive *in silico* analyses reveal specific changes in cytokine-nucleocapsid protein interactions, shedding light on potential modulators of the host immune response during infection. These findings offer valuable insights into the molecular mechanisms underlying viral pathogenesis and may guide the future development of targeted interventions. This manuscript serves as insight into the comparison of deep learning based AlphaFold2-Multimer and the semi-physicochemical based HADDOCK for protein-protein docking. We show the two methods are complementary in their predictive capabilities. We also introduce a novel algorithm for rapidly assessing the binding interface of protein-protein docks using graph edit distance: graph-based interface residue assessment function (GIRAF). The high-performance computational framework presented here will not only aid in accelerating the discovery of effective interventions against emerging viral threats, but extend to other applications of high throughput protein-protein screens.

## Introduction

SARS-CoV-2 evolution over the course of the pandemic has led to sustained, continued waves of infections. The variants of concern have shown a high degree of mutation relative to the prevailing strain at the time of their emergence. Most research has focused on the impact of mutations on the spike proteins’ ability to enter cells and evade antibodies, whether the antibodies are therapeutics or induced from vaccination or induced from prior infection (2, 3). The main reason for the focus on spike is driven by the fact that the most selective pressure on the virus is applied to the spike and the mutations have consequences for the therapeutics that are available. Outside of spike, the nucleocapsid (N) gene of SARS-CoV-2 has had lineage defining mutations in each of the variants of concern (4). A robust platform for studying the effects of nucleocapsid mutations on viral RNA delivery and expression has identified mutations in the Alpha and Delta variants that may contribute to higher transmissibility (5).

Another critical function of SARS-CoV-2 N is to bind and sequester a subset of 11 cytokines in order to disrupt immune signaling (6). The set of cytokines that SARS-CoV-2 binds is shared with another distantly related human betacoronavirus, HCoV-OC43, though HCoV-OC43 also binds to an additional 6 cytokines with high affinity (6, 7). Binding to at least one cytokine, CXCL12*β*, was determined experimentally for human betacoronaviruses SARS-CoV and MERS-CoV in addition to SARS-CoV-2 and HCoV-OC43. Binding of CXCL12*β* was shown to inhibit chemotaxis of macrophages in a transwell culture study. Other viruses also employ extracellular proteins that bind to specific subsets of cytokines with high affinity, most notably the SECRET-domain containing proteins of the poxviruses (8, 9). The presence or absence of certain SECRET-domain containing proteins has important consequences on the pathogenicity of the poxvirus.

An *in silico* docking screen of the panel of 11 cytokines tested experimentally against SARS-CoV-2 original reference strain and key variants of concern was developed in the current study. An example output of the docking analysis is shown in Figure 1 for CXCL12*β* with SARS-CoV-2 WA-1. The *in silico* screen was set up on a high-performance computing system. AlphaFold2-Multimer (10) and HADDOCK (11) were utilized in parallel to predict cytokine binding sites on N, as GDockScore (12) and PRODIGY (13) were used to further assess the properties of the predicted cytokine binding. The goals were to identify the cytokine binding sites of the experimental cytokine hits and determine if their binding has been impacted by continuing evolution in the human host over the course of the pandemic. N proteins from human betacoronaviruses and close relatives of SARS-CoV-2 were included in the *in silico* screen as well to test the conservation of cytokine binding in the broader betacoronavirus family. Finally, we wanted to test the ability of the *in silico* tools to identify interactions with cytokines using the full 64 cytokine panel utilized experimentally. The full cytokine panel docking screen could be utilized in order to track how evolution of N impacts this function continuing forward in the pandemic as well test other viral proteins for the ability to sequester cytokines.

**Fig. 1.**
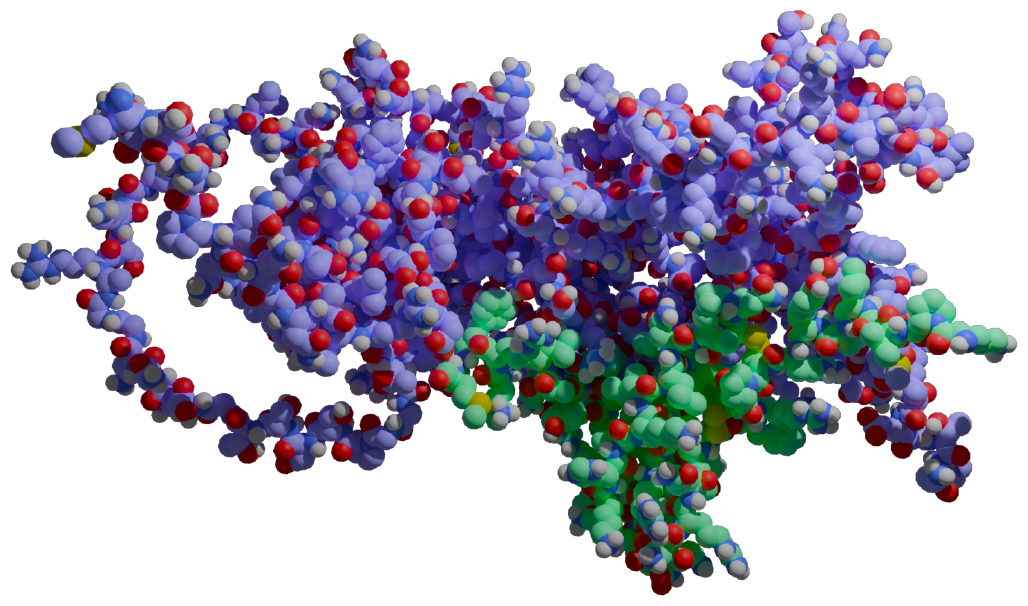
Human cytokine CXCL12*β* (in green) docked against the SARS-CoV-2 WA-1 nucleocapsid protein (in purple). Rendered with MolecularNodes (1).

## Results

We predicted 17 N protein structures across six betacoronaviruses (SARS-CoV, SARS-CoV-2, MERS-CoV, HCoV-OC43, RaTG13 and BANAL-20-52), all of which generally matched the expected topology of the N protein (14). For example, the predicted structure of the SARS-CoV-2 N WA1 strain had two structured domains that recapitulated two crystallized domains (PDB 6M3M for the N-terminal RNA binding domain, total RMSD 0.851Å and 6WZO for the dimerization domain, total RMSD 1.178Å) as well as three flexible domains with low AlphaFold2 confidence (pTM) that corresponded with the three intrinsically disordered domains. AlphaFold2 was also used to generate the structure of the 64 cytokines. The generated N and cytokine structures were used as inputs for HADDOCK, located here: https://github.com/tuplexyz/SARS-CoV-2_N-Cytokine_Docking/tree/main/haddock/inputs.

### Computing Benchmarks

Running on the MIT SuperCloud high-performance computing (HPC) environment (15), significant throughput was achieved for the numerous computational docking tasks.

### AlphaFold2 Performance

AlphaFold2-Multimer jobs took 493 ± 21 minutes of walltime per multimer run on nodes containing an Nvidia Volta V100 GPU, which contain 5,120 CUDA cores and 640 tensor cores. To complete the 1,088 AlphaFold2-Multimer experiments, 36 nodes were used, completing the entire set of experiments in approximately 12 days. A later subset was run on Ampere A100 GPUs to investigate if the newer generation GPU would increase speedups; the mean runtime decreased to 378 minutes ± 80 minutes (noting that the variability of duration increased). The A100 GPUs have more CUDA cores but fewer tensor cores, 6,912 and 432 respectively.

### HADDOCK Performance

HADDOCK jobs took 707 ± 40 minutes of walltime on Intel Xeon Phi 7210 cluster nodes, which contain 64 physical cores. On Intel Xeon Platinum 8260 nodes, which contain 48 physical cores, HADDOCK took 133 ± 92 minutes of walltime. The speedup on the Xeon Platinum nodes was likely due to the much newer CPU architecture and higher maximum clockspeed of the Xeon Platinum (3.9 GHz) compared to the Xeon Phi (1.5 GHz). While HADDOCK performed as quickly as 74 min on the Xeon Platinum nodes, it had a wide range of runtimes, depending on the cytokine and number of surface residues selected against which to dock. Cytokines with multiple chains (IL-12p70, IL-23, IL-27, and IL-35) took approximately three times as long. The 1,088 HADDOCK docking experiments were run on 64 of the Xeon Platinum 8260 nodes, completing the entire set of experiments in approximately 36 hours.

### Nucleocapsid Cytokine Binding Sites

HADDOCK and AlphaFold2-Multimer were used to identify potential binding sites of the cytokines with the nucleocapsid proteins (Figure 2). CXCL12*β* was chosen for initial representative structure modeling because it has been experimentally shown to bind to HCoV-OC43 and SARS-CoV-2 WA1 nucleocapsid in bio-layer interferometry assays (6, 7). SARS-CoV, MERS-CoV, HCoV-OC43 and SARS-CoV-2 nucleocapsid were additionally shown to inhibit CXCL12*β* dependent migration of macrophages in a transwell assay (6, 7). For the six betacoronaviruses (using SARS-CoV-2 WA1 as the representative strain of SARS-CoV-2), HADDOCK docked CXCL12*β* to locations around the dimerization interface, with contributing contacts from other domains of the protein. AlphaFold2-Multimer complexed the cytokine consistently at the dimerization interface, including interacting residues in the beta sheets of both the nucleocapsid dimerization interface and CXCL12*β*.

**Fig. 2.**
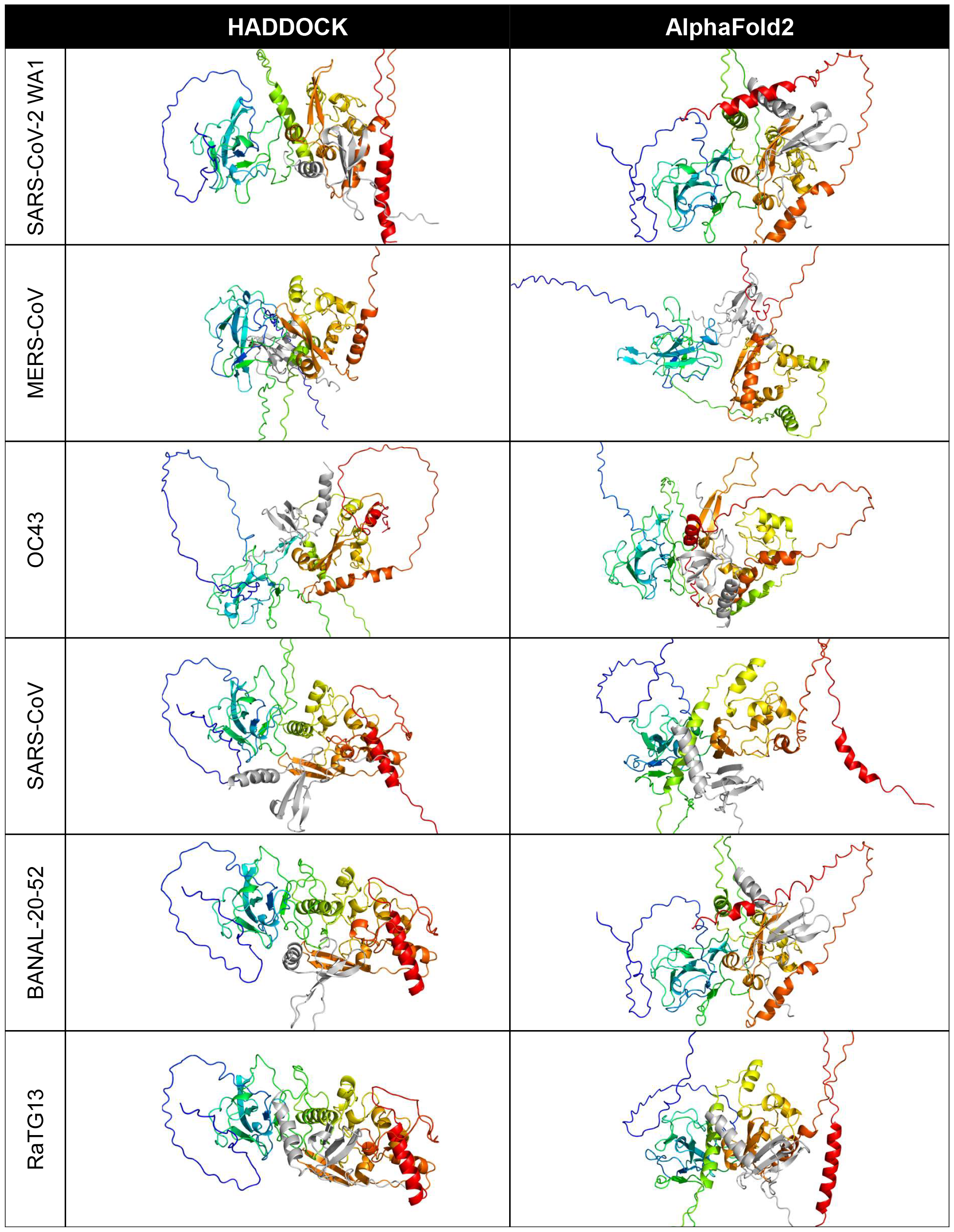
Predicted binding site of CXCL12*β* across selected coronaviruses. The proteins are shown in cartoon style with the N protein colored as rainbow (blue is the amino/N-terminus, red is the carboxyl/C-terminus) and the CXCL12*β* cytokine is colored in gray.

A closer look at the HADDOCK and AlphaFold2-Multimer docking of CXCL12*β* showed that HADDOCK did not include any beta sheet residues from the dimerization interface, but included polar contacts with residues on either side of the interface loop (S310, G335 and A336) (Figure 3). Other interacting residues were located in the structured N-terminal RNA binding domain and the flexible C-terminal region. The AlphaFold2-Multimer complex included two polar contacts of residues directly involved in dimerization (R319 and I320), as well as resides in the flexible C-terminal region. Two residues on CXCL12*β* involved in glycosaminoglycan (GAG) binding (H25 and R47) were involved in the HAD-DOCK predicted polar contacts, whereas the GAG-binding residues showed hydrogen bonding but no polar contacts in the AlphaFold2-Multimer predicted complex (16, 17). The SARS-CoV-2 WA1 nucleocapsid CXCL12*β* interaction was previously shown to be competitive with heparin sulfate and chondroitin sulfate (6).

**Fig. 3.**
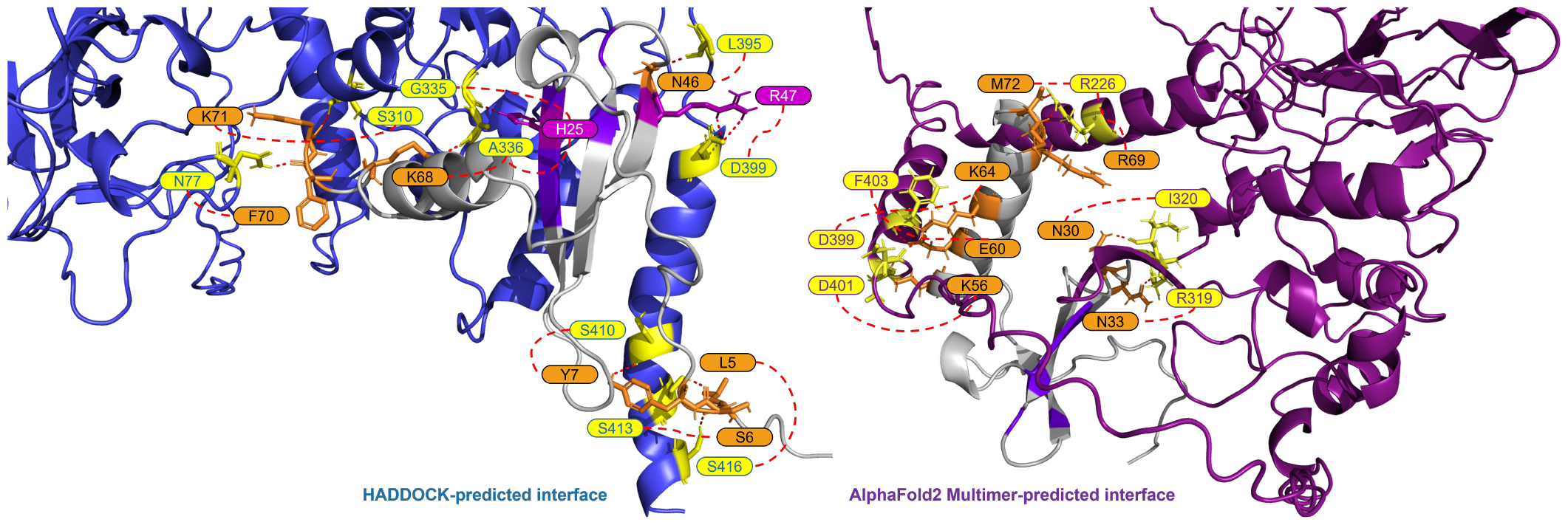
Predicted interface between the SARS-CoV-2 WA-1 N protein and CXCL12*β*. The proteins are shown in cartoon style with the N protein colored as periwinkle in the HADDOCK-predicted complex and dark magenta in the AlphaFold2 Multimer-predicted complex (with polar interacting residues shown as yellow sticks). The CXCL12*β* protein is shown in cartoon style in gray (with polar interacting residues shown as orange sticks). Residues in the GAG binding site on CXCL12*β* are shown in purple (with polar interacting residues shown as magenta sticks).

### Cytokine Strength

Across the 64 cytokines that were tested, binding affinities were highly correlated between the AlphaFold2-Multimer predictions and HADDOCK. However, the ranges of the scores may vary significantly. For example, the range of the HADDOCK van der Waals energies is [-109.82, -38.45] compared to [-56.76, -10.46] for the AlphaFold2-Multimer predictions (measured by FoldX). See Table 1.

**Table 1.**
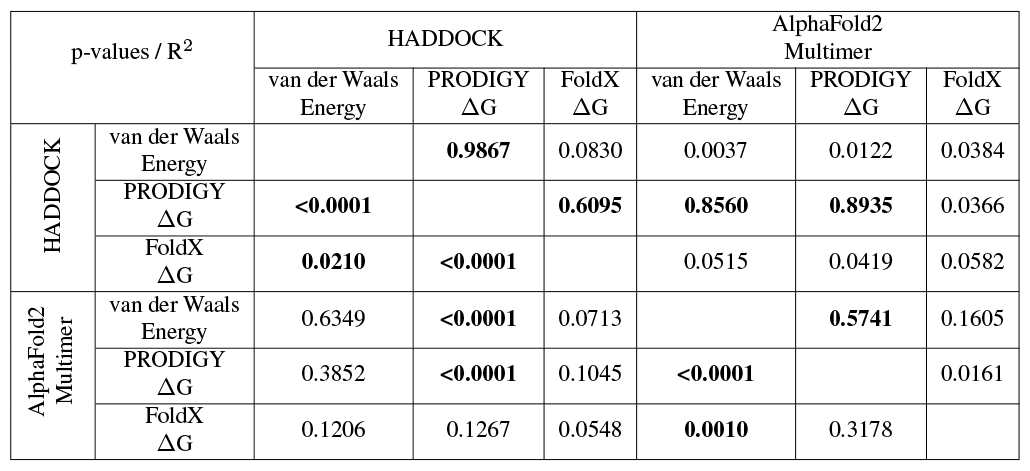
Correlations between AlphaFold2-Multimer and HADDOCK docking predictions across various binding metrics. Significant correlations are shown in **bold**.

### Genetic Distance

Correlating the cytokine binding affinity metrics with Fitch distance of each SARS-CoV-2 variants’ N protein from the SARS-CoV-2 WA1 N protein, there were significant positive correlations for some of the 64 cytokines, including some which were hits in experimental SARS-CoV-2 and/or HCoV-OC43 screens (6, 7). Namely, CCL28, CXCL10, CXCL11, CXCL13 and CXCL14 from the HAD-DOCK predictions show that the binding Gibbs energy predicted by PRODIGY weakened as the genetic distance of the N protein increases from SARS-CoV-2 WA1 (Figure 4 H.a). No significant trends were observed for the predicted AlphaFold2 Gibbs energy by PRODIGY of the cytokines from the experimental screens (Figure 4 AF.a). When correlating the Fitch distance to predicted Gibbs energy by FoldX, only CXCL12*α* showed a significant correlation of the experimental hits. For this cytokine, both HADDOCK and AlphaFold2 predictions yield a negative correlation with genetic distance, indicating a stronger association with CXCL12*α* over the course of pandemic (Figure 4 H.b and AF.b). The correlation between Fitch distance van der Waals energy was significant for only CCL28 of the experimental hits, again a positive correlation predicted by HADDOCK indicating less favorable binding (Figure 4 H.C and AF.c). Several cytokines not identified in either experimental screen had significant positive correlations with Fitch distance. However, cytokines that were non-binders in the screen and had increased Gibbs energy or van der Waals energy would still be expected to be non-binders. Nine cytokines that were negative in both screens had a significant negative correlation in one of the three metrics of binding strength: CXCL8, CCL27, IFN*λ*1, TNF*α*, CCL2, CCL3, IL-6, IL6R*α* and IL-18BP. Experimental validation could verify if any of these cytokines are inhibited variants of SARS-CoV-2 N even though SARS-CoV-2 WA1 N did not inhibit them.

**Fig. 4.**
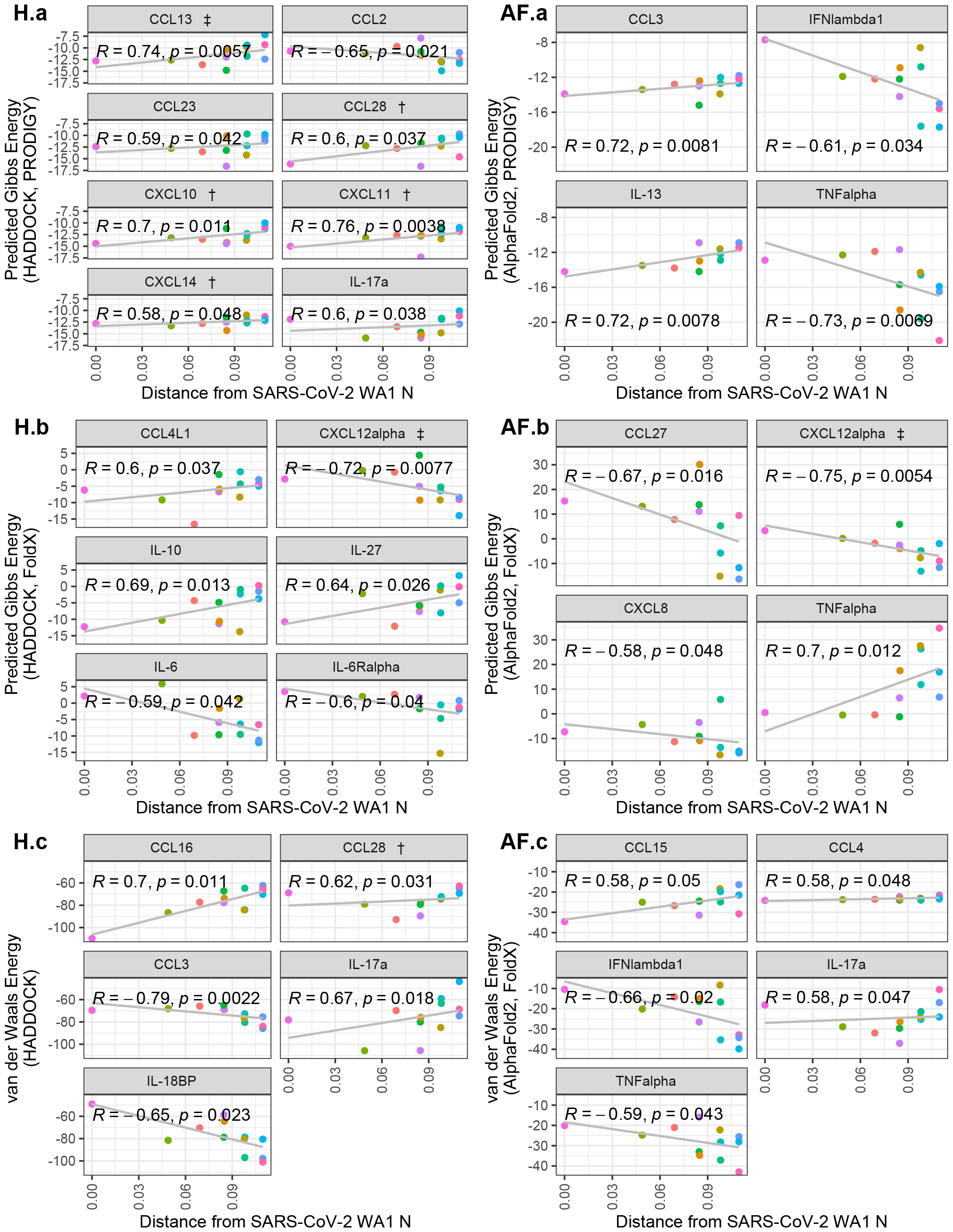
Scatterplots of cytokine binding affinity for all cytokines with a statistically-significant correlation to genetic distance. Shown is the relationship between the Fitch distance from the SARS-CoV-2 WA1 N variant vs. HADDOCK and AlphaFold2 predicted binding affinity metrics. Each point represents a SARS-CoV-2 variant, with the most distant points representing Omicron subvariants. Cytokines referenced in López-Muñoz et al. (2022) are denoted by † and ‡ for WA-1/OC43 and OC43 only hits, respectively.

A phylogenetic tree using only N protein sequences was generated using RAxML (Figure 5). The expected phylogenetic topology was observed of a tree generated using both reference coronaviruses like HCoV-OC43, SARS-CoV, and MERS-CoV along with SARS-CoV-2 variants. SARS-CoV-2 variants were grouped in a monophyletic clade for this analysis.

**Fig. 5.**
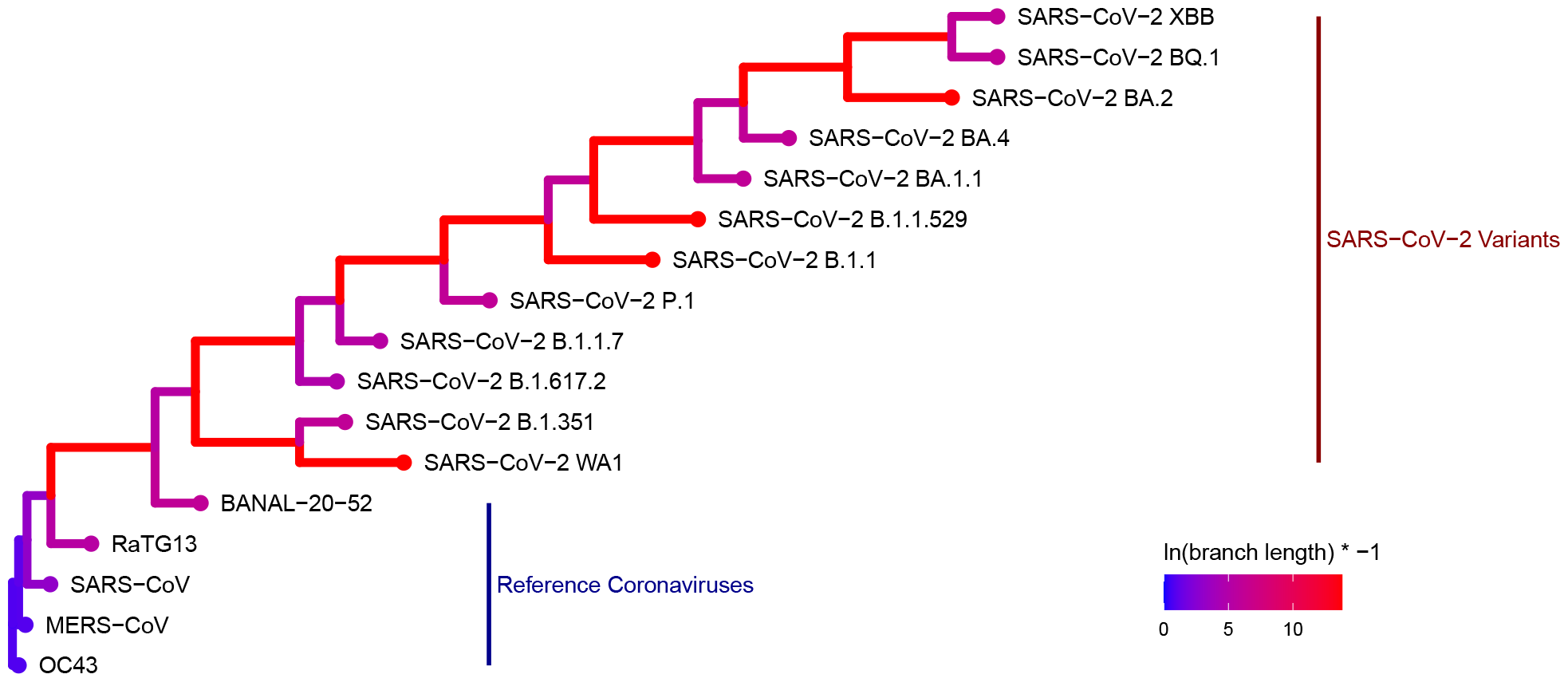
Phylogenetic tree generated with RAxML depicting the genetic distance of the 17 coronavirus N proteins used in this study. (Colors represent the log-transformed branch length: *ln*(branch length) *× −*1.)

### Breadth of CXCL12*β* Binding

Graph Edit Distance (GED) provided a view into how the N-protein receptor (i.e., the binding pocket) to CXCL12*β* changed between different N-proteins. A novel algorithm was developed in order to determine GED of the bind site, named graph-based interface residue assessment function (GIRAF). Both AlphaFold2 and HADDOCK structures concurred with relatively low GED of SARS-CoV-2-B.1.1 and SARS-CoV-2-B.1.617.2-DeltaA (Figures 6A, 6B, respectively). AlphaFold2 had high GED for MERS-CoV and SARS-CoV-2-BQ.1, suggesting far evolutionary distance in the binding pocket compared to the SARS-COV2-WA1 baseline (Figure 6A).

**Fig. 6.**
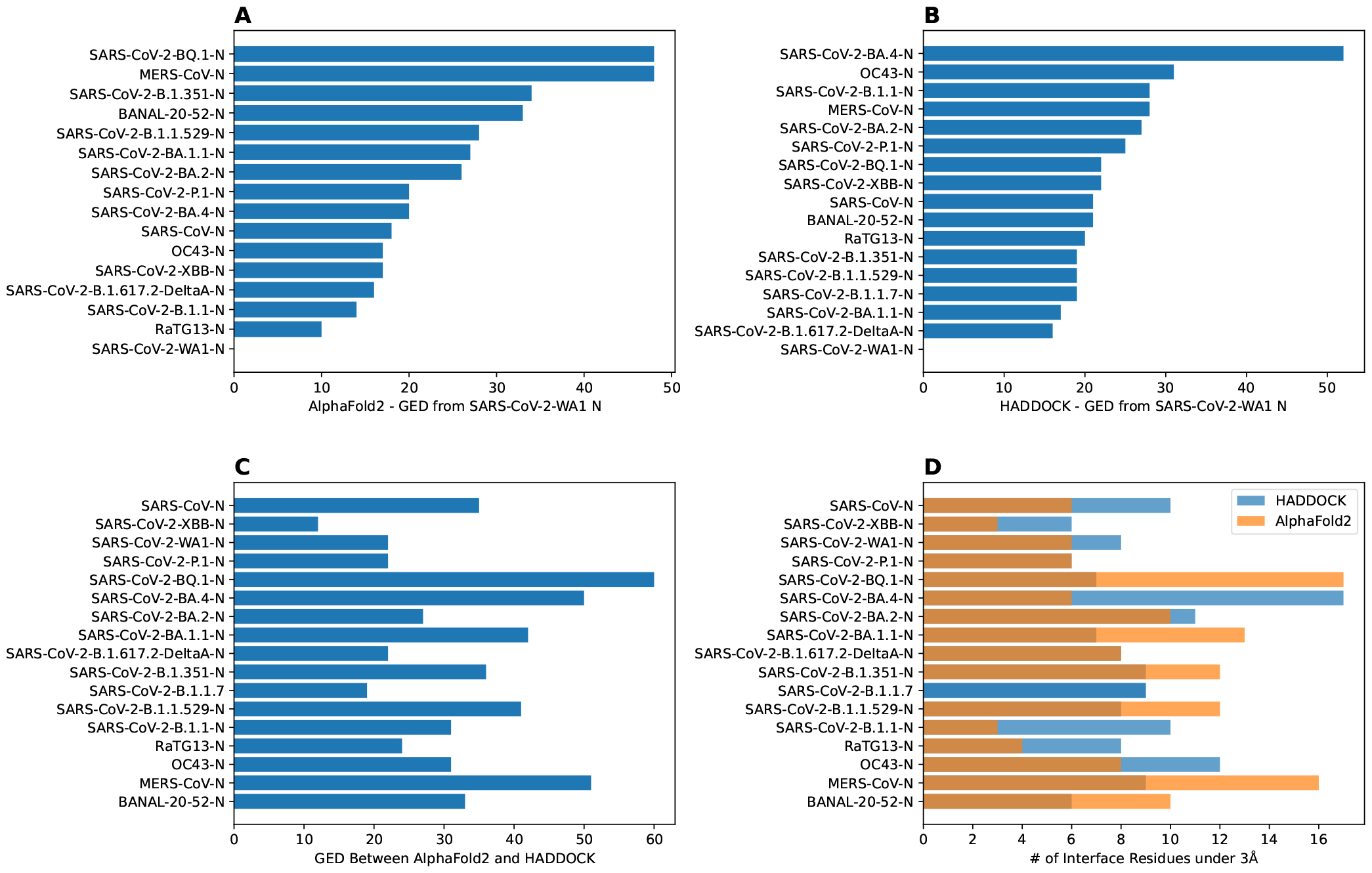
Bar charts depicting the graph edit distance for CXCL12*β* using GIRAF. A) AlphaFold2 GED from SARS-CoV-2 WA1 N. Note, there were no interface residues less than 3Å for SARS-CoV-2-B.1.1.7. B) HADDOCK GED from SARS-CoV-2 WA1 N. C) GED between AlphaFold2 and HADDOCK SARS-CoV-2 WA1 N. D) Number of interface residues under 3Å.

There was marked disagreement between AlphaFold2 and HADDOCK for SARS-CoV-2-BQ.1 (Figure 6C). This was partly due to the discrepancy in the number of interfacing residues on the AlphaFold2 structure compared to the HAD-DOCK structure (Figure 6D). AlphaFold2 and HADDOCK reported a similar number of interfacing residues on other N proteins; both agreed the number of interfacing residues on SARS-CoV-2-XBB and SARS-CoV-2-P.1 as relatively low. AlphaFold2 reported six residues on SARS-CoV-2-WA1 compared to HADDOCK’s eight residues.

## Discussion

*In silico* structural prediction and molecular docking tools were utilized in order to interrogate the potential evolutionary change of the interaction between SARS-CoV-2 nucleocapsid proteins and 64 human cytokines. Other betacoronaviruses were also included in the analysis, as several human betacoronaviruses had previously been shown to have a similar cytokine inhibitory phenotype. Additionally, HCoV-OC43 was shown to bind 17 human cytokines, including all 11 cytokines that SARS-CoV-2 bound despite being distantly related (6, 7). We identified the nucleocapsid dimerization domain as an important site of multiple cytokine interactions, with AlphaFold2 consistently identifying the dimerization loop specifically as the interaction site. We also identified five cytokines from the experimental screens (CCL28, CXCL10, CXCL11, CXCL13, CXCL14) that had significantly reduced binding to N as SARS-CoV-2 evolved by at least one *in silico* metric, and only one cytokine from the experimental screens that had increased binding (CXCL12*α*). Interestingly, only one of the five cytokines with reduced binding was from the HCoV-OC43 only, but the only cytokine with increased binding, CXCL12*α*, was from the HCoV-OC43 screen only. Finally, we identified nine cytokines that were negative in both experimental screens but had increased interactions with N variants. These cytokines could be potential new targets of inhibition by N.

We expected that interactions with N and cytokines would generally get stronger as SARS-CoV-2 co-evolved with humans. Contrary to our hypothesis, we found that the many statistically significant changes were weakening of the cytokine interaction as mutations accumulated. Multiple variants of the Omicron group were included in the analysis and were the most distant from the reference WA1 strain. The decreasing capacity for inhibition of cytokine signaling could be at least partially responsible for the decreased severity of disease observed with Omicron. Of the four cytokine that HADDOCK predicted would bind with decreased affinity, CXCL10 stands out as it is a strong predictor of disease severity (18–20). Decreased disregulation of CXCL10 by Omicron N could contribute to any related phenotypic change towards lower severity.

In addition to interrogating how the binding might change over the course of the pandemic for the cytokines that were experimentally shown to bind to SARS-CoV-2 WA1 N, we also identified a series of cytokines that may be inhibited by Omicron that were not inhibited by the original strain of SARS-CoV-2. These include predicted increased binding of CXCL8, CCL27, IFN*λ*1, and TNF*α* by AlphaFold2 and CCL2, CCL3, IL-6, IL6R*α* and IL-18BP by HADDOCK. Though the predicted affinity and/or van der Waals associated is lower for the variants compared to the reference strain, it is not certain that these cytokines would be inhibited because it is unclear if the lower affinities translate to physiological binding and the predicted binding sites may not be conducive to inhibition of cytokine function.

The predicted SARS-CoV-2 N binding to cytokine shares several features with the better characterized SECRET-cytokine binding. The binding to N typically involves a beta sheet to beta sheet interaction. One of the two main beta sheets involved in cytokine binding is the dimerization interface of SARS-CoV-2 N. The beta sheet interaction frequently involves the oligomerization interface of the cytokine. The binding both the GAG-binding domain oligomerization interface stabilized by a flexible arm describe the interaction motif of the SECRET proteins, at least one interaction of which has been crystallized (PDB 3ONA) (9).

Computational approaches to modeling binding are an attractive solution for variant tracking and biosurveillance in order to deal with the huge influx of sequences. Utilization of structural and docking predictions could be used to test the functional interactions as new variant sequences are identified. Previous computational studies have provided accurate early insights into viral mutations and their affects on human health (21–25).

In this work, we have not only showcased the utility of *in silico*-based protein modeling, but also the importance of scalability through high-performance computing. HPC-enabled frameworks, as used here, allow for tremendous throughput improvement, reducing costly and slow lab-based workloads. This approach will be important for proactively determining when a pathogen has acquired new phenotypes, such as increased transmissibility, pathogenicity, or zoonotic potential. In the long term, robust computational assays for several functions would need to be developed in order to track those higher-order viral features such as transmissibility.

## Methods

### Overall Approach

The docking experiment set in this study contained 64 human cytokines (supplemental table) and the nucleocapsid (N) protein from 17 different coronaviruses, 12 of which were from SARS-CoV-2 variants (Table 2). The experiment generation consisted of all possible combinations of the cytokines and N proteins for a total of 1,088 cytokine-N protein complexes.

**Table 2.** Coronavirus strains and NCBI accession numbers.

| Virus | Strain | Accession |
| --- | --- | --- |
| BatCoV | BANAL-20-52 | MZ937000.1 |
| MERS-CoV | England1 | NC_038294.1 |
| HCoV | OC43 | NC_006213.1 |
| BatCoV | RaTG13 | MN996532.2 |
| SARS-CoV | BJ01 | AY278488.2 |
| SARS-CoV-2 | B.1.1 | MT233522.1 |
| SARS-CoV-2 | B.1.1.529 | OL919777.1 |
| SARS-CoV-2 | B.1.1.7 | OP879258.1 |
| SARS-CoV-2 | B.1.351 | MW571126.1 |
| SARS-CoV-2 | B.1.617.2 | MW931310.1 |
| SARS-CoV-2 | BA.1.1 | OP969456. |
| SARS-CoV-2 | BA.2 | OP968961.1 |
| SARS-CoV-2 | BA.4 | ON324341.1 |
| SARS-CoV-2 | BQ.1 | OP839317.1 |
| SARS-CoV-2 | P.1 | MW520923.1 |
| SARS-CoV-2 | XBB | OP847716.1 |

Each experiment was submitted through both AlphaFold2-Multimer and HADDOCK docking systems. These tools generated numerous PDB files of predicted protein complex conformations. From these outputs, the best representative complex PDB structure was selected through various filtering techniques (described below) and then compared across the full experiment set. See Figure 7.

**Fig. 7.**
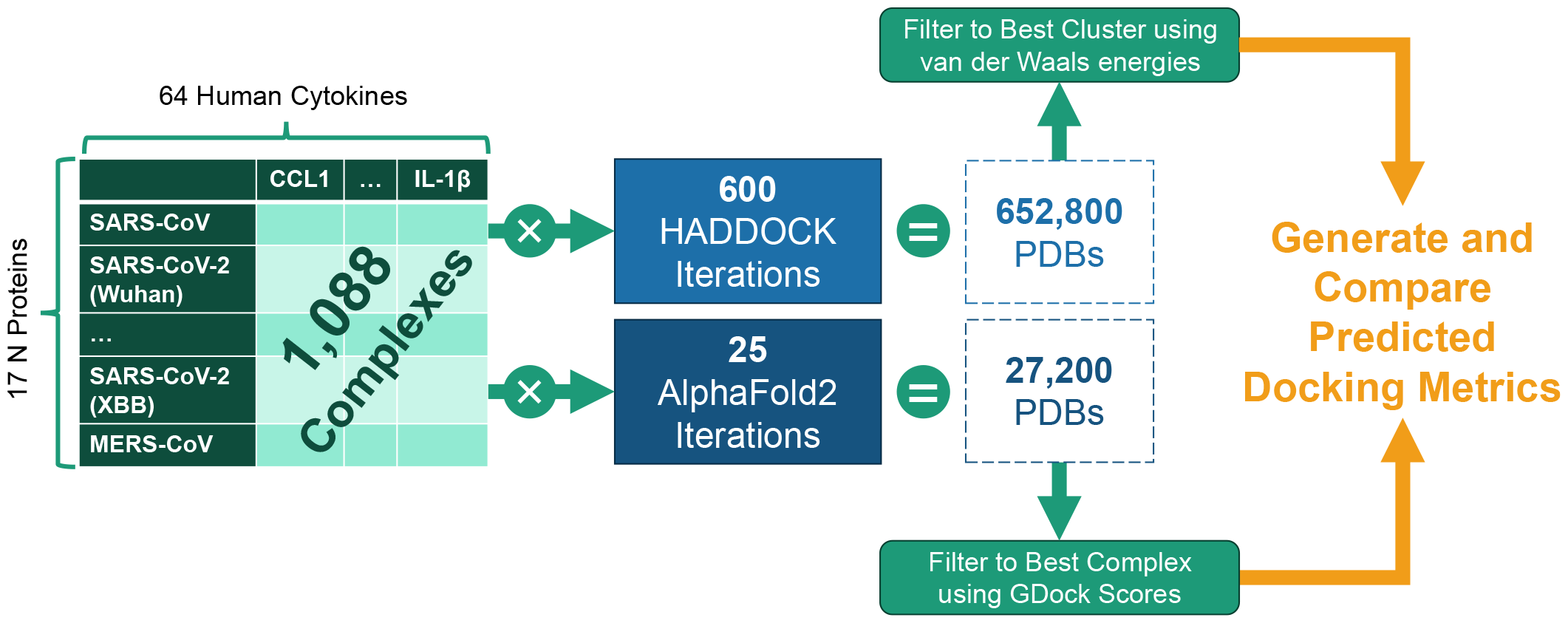
Experiment generation workflow for the 1,088 cytokine-N complexes.

### AlphaFold2 Protein Complex Prediction

AlphaFold2 (version 2.3.2) was compiled into a Singularity container and queried against a reference database constructed on 2023-04-12 according to the author’s instructions (26). The database was compressed into a SquashFS file and bound directly to the Singularity container. The specification for the AlphaFold2 Singularity container can be found at: https://github.com/mit-ll/AlphaFold.

AlphaFold2 was run in multimer mode and used default settings to construct 5 randomly seeded models with 5 predictions each, for a total of 25 models (PDBs) per complex. Amber relaxation was used on all resulting 25 models. AlphaFold2 was run on both standalone and high performance computing platforms.

For the standalone system, we used an NVIDIA DGX A100 80GB server. The system was equipped with dual AMD Rome 7742 CPUs, 2TB RAM, and 8 NVIDIA A100-SXM 80GB GPUs.

The MIT Lincoln Laboratory TX-GAIA and MIT SuperCloud (15) systems were the environments used for the HPC-enabled AlphaFold2 pipeline prototyping and performance benchmarking. The compute nodes each consisted of a Intel Xeon Gold 6248 2.5 GHz CPU with 40 cores, 377GB RAM, and Intel Omni-Path with 2 NVIDIA Tesla V100 GPUs.

Signal peptides were identified and removed from cytokine sequences using SignalP6.0 web portal^1^ before structure prediction with AlphaFold2 (27).

### AlphaFold2 Confidence

AlphaFold2 confidence was given as the interface predicted template model score plus predicted template model score (ipTM+pTM). The ipTM was weighted by 80% and the pTM was weighted by 20%, as described by the authors (10). The total AlphaFold2 confidence (ipTM+pTM) ranged from [0, 1], where 1 was the highest confidence.

### GDock Score

*GDockScore*, a graph-based deep learning model to assess the docking of two proteins, was included to evaluate AlphaFold2 models (12). The model was pretrained by the original authors on docking outputs generated from Protein Data Bank, RosettaDock, HADDOCK decoys, and the ZDOCK Protein Docking Benchmark- to include a wide variety of protein complexes and ensure generalization. *GDockScore* achieved state-of-the-art on the CAPRI Score Set, a challenging dataset for developing docking scoring functions (28). *GDockScore* ranged from 0 to 1, where a score of 0 coincided to an unfeasible complex and a score of 1 coincided with high probability that the protein complex is comparable to a high quality CAPRI complex. The authors showed a *GDockScore* of under 0.2 approximates to a complex that was unlikely to exist (i.e., bad protein-protein dock) and a *GDockScore* range from 0.2 to 1 coincided with an increasing predicted quality.

### Best AlphaFold2 Structure Selection

AlphaFold2 ranked the 25 structures by confidence (AF2 conf.). We added the raw AF2 conf. to the GDock score (i.e., total score) for all 25 structures and selected the highest score for each multimer experiment. The highest total score was not necessarily the best ranked from AF2 conf. alone. The best 1,088 structures were used for all downstream analysis. See Figure 8

**Fig. 8.**
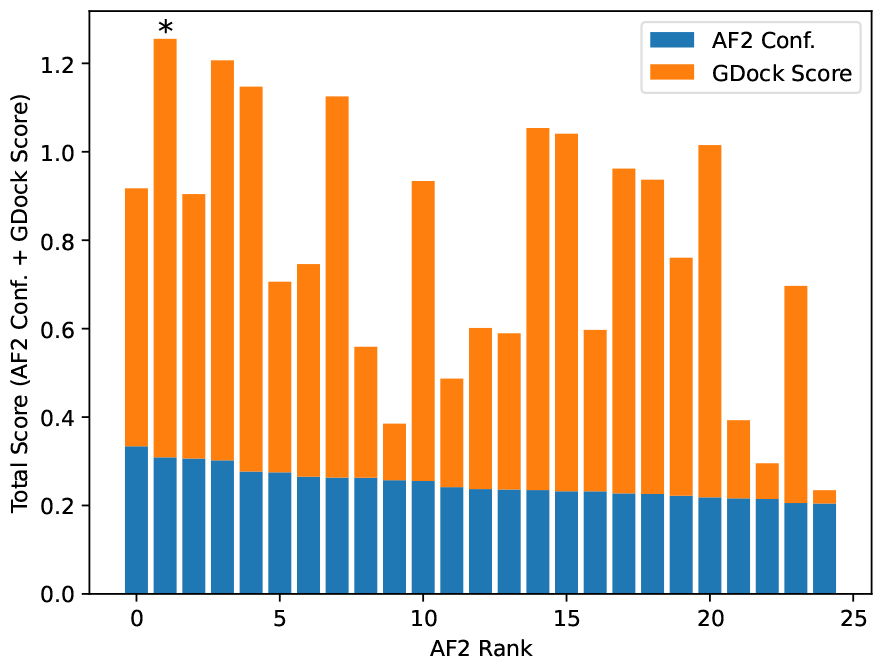
Example total score calculation for SARS-CoV-2-WA1 bound to CXCL12*β*. AlphaFold2 confidence was calculated by ipTM+pTM (see text for details). The AlphaFold2 rank 0 had a slightly higher confidence than the rank 1, however the rank 1 structure had a significantly higher GDock score which led to a higher total score. In this case, the rank 1 structure was selected as the best structure.

### HADDOCK Docking

For each of the 1,088 experiments generated in this study, HADDOCK v2.4 (11) was used to dock the N protein and cytokine in each experiment.

HADDOCK, High Ambiguity Driven protein-protein DOCKing, is a biomolecular modeling software that provides docking predictions for provided structures using an information-driven flexible docking approach.

For this study, we utilized a Docker containerized version of HADDOCK, which contains all of the software dependencies to allow HADDOCK to run more readily in an HPC environment. HADDOCK was run on both 64 physical cores (Intel Xeon Phi 7210) and 48 physical cores (Intel Xeon Platinum 8260). HADDOCK needed to be compiled against the number of physical cores; more information can be found at: https://github.com/colbyford/HADDOCKer, or on Dock-erHub at: https://hub.docker.com/r/cford38/haddock.

### HADDOCK Experiment Setup

Run parameters for each experiment were generated programmatically, defining the N protein as the static object around which the cytokine protein is positioned and measured. Other experiment files were also copied or created programmatically that include the scripts to run the docking process, define restraints, and specify input PDB files.

HADDOCK required the definition of active/inactive residue restraints (AIRs) to help guide the protein docking process. To avoid biasing the docking placement of the cytokine, all surface residues were selected as “active” and were then included in the AIR file on which to dock.

The logic for this programmatic generation of HADDOCK experiment files is available in the supplementary GitHub repository.

### HADDOCK Outputs

The HADDOCK docking process consisted of three steps:

*it0*: Randomization of orientations and rigid-body minimization

*it1*: Semi-flexible simulated annealing through molecular dynamics simulations in torsion angle space

*itw*: Refinement by energy minimization in Cartesian space with explicit solvent (i.e., in water)

Each of the above steps generated 200 PDB files of the proteins in a complex for a total of 600 PDB files. From the final, water-refined (*itw*) set of outputs, the “best” PDB file was retrieved from the cluster of predictions with the lowest van der Waals energy. This representative PDB of each Ncytokine complex was then used in subsequent analyses and comparisons.

### HADDOCK Metrics

The HADDOCK system outputted multiple metrics for the predicted binding affinities and an output set of PDB files containing the N protein docked against the cytokine protein. Some main metrics included:

- van der Waals intermolecular energy (*Evdw*)
- Electrostatic intermolecular energy (*Eelec*)
- Desolvation energy (*Edesol*)
- Restraints violation energy (*Eair*)
- Buried surface area (*BSA*)
- HADDOCK score: 1.0*Evdw* + 0.2*Eelec* + 1.0*Edesol* + 0.1*Eair*

### PRODIGY ΔG Prediction

PRODIGY, a tool to predict the binding affinity of protein-protein complexes from structural data, was used on each complex in this study (13). The predicted binding affinities were reported as Gibbs energy, shown as ΔG (in Kcal/mol units).

The predicted ΔG values were calculated by counting the number of various interfacial contacts (ICs) between the chains of the input complex (N protein and cytokine) along with some properties of the non-interacting surfaces (NIS) using the following equation:

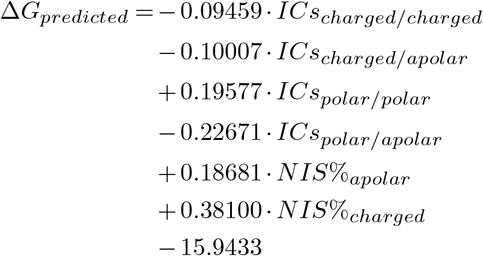

### FoldX ΔG Prediction

FoldX (29), a tool that evaluates protein-protein complex interactions and their stability, was also used to predict ΔG for all of the complexes. In the command line, the AF2 and HADDOCK structures were first repaired using the PDB repairing function, RepairPDB, which minimized the complex energy by rotating identified side chain residues.

FoldX calculated ΔG for the repaired structures using a linear combination of empirically derived energy terms including van der Waals, solvation energy for apolar and polar groups, and electrostatic contribution of charged groups.

### Graph-based Interaction Residue Assessment Function (GIRAF)

The interfaces of all complexes were first processed with *INTERCAAT* (30) to produce a full list of all interacting residue pairs between the N-protein and bound cytokine. A bigraph was created using Python NetworkX v3.1 (31) where the nodes were defined as each residue and the edges were defined as the predicted distance between the residues (in angstroms) by *INTERCAAT*. Distances longer than 3 angstroms were discarded. A lookup table was constructed for the 17 N-proteins using a consensus sequence from multiple sequence alignment (Geneious, Dotmatics, Inc) so that each residue number was aligned with the consensus. Baseline graphs were generated for SARS-CoV-2-WA1 with CXCL12*β* from both AlphaFold2 and HAD-DOCK predicted complexes. Graphs were then generated for the other N-proteins with CXCL12*β*. Graph Edit Distance (GED) was computed between each of the other N-proteins with SARS-CoV-2-WA1. GED was then computed between each N:CXCL12*β* complex from AlphaFold2 and HADDOCK.

The number of interfacing residues on N-protein from both AlphaFold2 and HADDOCK were enumerated. GED between a pair of graphs was defined as follows:

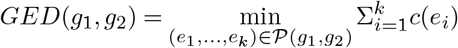

GED minimization was capped at 10 seconds as we saw that further significant cost minimization over the set of edit paths did not occur beyond this point. GED costs were equally weighted a value of 1 for substitutions, insertions, or deletions.

Example bigraphs are shown in Figure 9 where interfacing residues between the N protein and a cytokine, shown as labeled dots, are connected to one another.

**Fig. 9.**
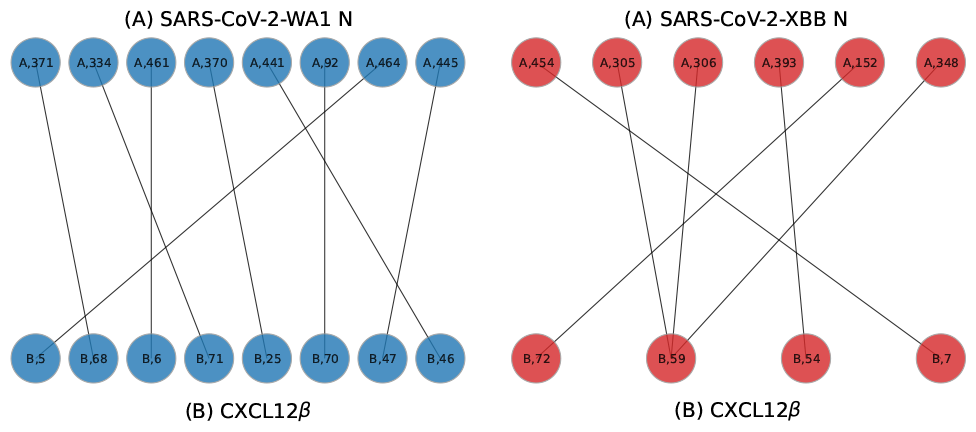
Example bigraphs for SARS-CoV-2-WA1 (A chain, left) and SARS-CoV-2-XBB (A chain, right) bound to CXCL12*β* (B chains) on models predicted by HAD-DOCK.

### Protein Structure and Results Visualization

Protein complexes were visualized using using PyMOL (32), except Figure 1. PyMol’s analysis capabilities were employed to detect and show interfacing residues (polar interactions within 3.0Å) between the N protein and cytokine in each complex. Graphs were rendered using the ggplot2 v3.4.2 R package (33) or Matplotlib v3.8.1 (34). The phylogenetic tree figure was rendered using the ggtree v3.6.2 R package (35).

### N Protein Alignment, Genetic Distance, and Tree Generation

The sequences for the N protein of each variant were aligned using Muscle v3.8.425 (36) and was written in FASTA format. From the aligned FASTA file, the pairwise distances were computed using the Fitch matrix (37) from the seqinr v4.2.30 R package (38). For Figure 4, the distance from the SARS-CoV-2 WA1 isolate was used as the reference point.

A maximum likelihood phylogenetic tree of the N protein alignment was generated with RAxML v8.2.12 (39). This tree was generated using the “rapid bootstrap” mode with 100 replicates and rooted on the OC43 taxon.

## Author Contributions

PJT and RJ3 conceptualized and obtained funding for the project. PJT and CTF drafted the original manuscript draft. RJ3 and AMM developed the pipeline to run AlphaFold2 and provided assistance with debugging the HADDOCK software container. CTF developed the pipeline to run HAD-DOCK and PRODIGY, generated the PDB model figures, performed correlations for cytokine binding affinity and Fitch distance, and the phylogenetic tree. RJ3 developed the GDockScore pipeline to select the AlphaFold2 structures and Graph Edit Distance (G-RAF) algorithm. AEM performed FoldX and statistical analysis. PJT, CTF, RJ3, and AEM all analyzed data and edited the manuscript.

## Acknowledgements

The authors would like to thank Alberto D. López-Muñoz and Jonathan Yewdell at the National Institutes of Health for guidance on understanding coronavirus nucleocapsid and cytokine interactions. The authors also want to acknowledge Darrell Ricke (MIT LL) for initial help with debugging AlphaFold2, Keegan Quiqley (MIT LL) and Menghua Rachel Wu (MIT CSAIL) for helpful suggestions, the MIT SuperCloud and Lincoln Laboratory Supercomputing Center for providing resources and consultation, the MIT BeaverWorks facilities, and Jon Halter from the UNC Charlotte University Research Computing group.

## Data availability statement

All code, data, results, and additional analyses are openly available on GitHub at: https://github.com/tuplexyz/SARS-CoV-2_N-Cytokine_Docking.

This repository includes the PDB files for the top 1,088 complexes for both AlphaFold2 and HADDOCK experiments, all output metrics, experiment generation and data preparation logic, HPC submission scripts, and the code for and post-processing analyses and generating figures.

## Legal and funding statement

DISTRIBUTION STATEMENT A. Approved for public release. Distribution is unlimited.

This material is based upon work supported by the Department of the Air Force under Air Force Contract No. FA8702-15-D-0001. Any opinions, findings, conclusions or recommendations expressed in this material are those of the author(s) and do not necessarily reflect the views of the Department of the Air Force.

## Supplementary Material

**Table S1.** Cytokines along with their NCBI accession IDs.

| Cytokine | Family | Accession | Cytokine | Family | Accession |
| --- | --- | --- | --- | --- | --- |
| CCL1 | β-chemokine | NP_002972.1 | IL-1α | Interleukin | AAC41702.1 |
| CCL2 | β-chemokine | NP_002973.1 | IL-1β | Interleukin | NP_000610.2 |
| CCL3 | β-chemokine | NP_002974.1 | IL-6 | Interleukin | NP_742152.1 |
| CCL3L1 | β-chemokine | NP_066286.1 | IL-6Rα | Interleukin | KAI2552237.1 |
| CCL4 | β-chemokine | NP_002975.1 | IL-10 | Interleukin | CAA27448.1 |
| CCL4L1 | β-chemokine | NP_996890.1 | IL-12p70 | Interleukin | NP_000567.1 |
| CCL5 | β-chemokine | NP_002976.2 | IL-13 | Interleukin | NP_000591.1 |
| CCL7 | β-chemokine | NP_006264.2 | IL-17a | Interleukin | NP_000556.1 |
| CCL8 | β-chemokine | NP_005614.2 | IL-18 | Interleukin | NP_000563.1 |
| CCL11 | β-chemokine | NP_002977.1 | IL-18BP-Fc | Interleukin | NP_000873.2; NP_002178.2 |
| CCL13 | β-chemokine | NP_005399.1 | IL-23 | Interleukin | NP_002179.2 |
| CCL14 | β-chemokine | NP_116739.1 | IL-27 | Interleukin | NP_002181.1 |
| CCL15 | β-chemokine | NP_116741.2 | IL-35 | Interleukin | NP_001373349.1 |
| CCL16 | β-chemokine | NP_004581.1 | TNF-α | Tumor Necrosis Factor | NP_001034749.1 |
| CCL17 | β-chemokine | NP_002978.1 | TNF-β | Tumor Necrosis Factor | NP_057668.1; 3D87_B |
| CCL18 | β-chemokine | NP_002979.1 | IFN-β | Interferon | NP_663634.2; Q14213.2 |
| CCL19 | β-chemokine | NP_006265.1 | IFN-γ | Interferon | NP_001341512.1; NP_005746.2 |
| CCL20 | β-chemokine | NP_004582.1 | IFN-λ1 | Interferon | NP_000585.2 |
| CCL21 | β-chemokine | NP_002980.1 | IFN-ω | Interferon | NP_001153212.1 |
| CCL22 | β-chemokine | NP_002981.2 | IFN-α2 | Interferon | NP_002986.1 |
| CCL23 | β-chemokine | NP_005055.3 | | | |
| CCL24 | β-chemokine | NP_002982.2 | | | |
| CCL25 | β-chemokine | NP_005615.2 | | | |
| CCL26 | β-chemokine | NP_001358867.1 | | | |
| CCL27 | β-chemokine | NP_006655.1 | | | |
| CCL28 | β-chemokine | NP_683513.1 | | | |
| CXCL1 | α-chemokine | CAG33707.1 | | | |
| CXCL2 | α-chemokine | NP_001502.1 | | | |
| CXCL3 | α-chemokine | NP_002080.1 | | | |
| CXCL4 | α-chemokine | NP_002081.2 | | | |
| CXCL5 | α-chemokine | NP_002610.1 | | | |
| CXCL6 | α-chemokine | NP_002985.1 | | | |
| CXCL7 | α-chemokine | NP_002984.1 | | | |
| CXCL8 | α-chemokine | NP_002695.1 | | | |
| CXCL9 | α-chemokine | NP_000575.1 | | | |
| CXCL10 | α-chemokine | NP_002407.1 | | | |
| CXCL11 | α-chemokine | NP_001556.2 | | | |
| CXCL12α | α-chemokine | NP_005400.1 | | | |
| CXCL12β | α-chemokine | NP_954637.1 | | | |
| CXCL13 | α-chemokine | NP_000600.1 | | | |
| CXCL14 | α-chemokine | NP_001358487.1 | | | |
| CXCL16 | α-chemokine | NP_004878.3 | | | |
| XCL1 | γ-chemokine | NP_001373738.1 | | | |
| CX3CL1 | δ-chemokine | NP_000596.2 | | | |

